# AQP1- A regulatory factor associated with brown adipose tissue-silencing

**DOI:** 10.1101/2024.09.23.614599

**Authors:** Chloe M Cheng, Christopher J Blay, Pei-Yin Tsai, Muying Li, Kaydine Edwards, Yue Qu, Yang Liu, Nina Buettner, Claire Walter, Mary Snyder, Ines PD Costa, Olivier Devuyst, Joeva J Barrow

## Abstract

The activation of non-shivering thermogenesis (NST) in brown adipose tissue (BAT) by environmental cold challenge yields strong metabolic benefit in the face of diet-induced obesity (DIO). Yet, a critical barrier to leveraging brown fat NST for therapeutic use against metabolic disease is that BAT is silenced and inactive at physiological ambient temperature conditions in humans. The mechanisms that govern this silencing process remain poorly understood. Here, we identified a putative BAT-silencing factor, aquaporin-1 (AQP1), in brown fat from wild-type (WT) mice via proteomics analysis. We generated the first BAT-specific AQP1 knockout mice (AQP1-KO) and revealed that AQP1-KO could activate NST under BAT silencing environmental conditions and that the AQP1-KO mice were significantly protected against DIO and metabolic dysfunction compared to Flox controls. We found that AQP1-KO mice on high fat diet (HFD) had reduced weight gain through reductions in fat mass, improved glucose tolerance, and increased whole body energy expenditure compared to Flox control mice. Mechanistically, we show that AQP1 ablation in mice had upregulated gene expression related to the electron transport chain (ETC) and mitochondrial translation contributing to the activation of NST under BAT environmental silenced conditions.

**Significance Statement:** Novel strategies to combat obesity-associated metabolic dysfunction are urgently needed to curb the growing obesity epidemic. Investigation of brown adipose tissue (BAT) silencing mechanisms may reveal novel therapeutic targets that when ablated, can activate BAT to increase energy expenditure and protect subjects against the metabolic dysfunction associated with obesity. We have identified Aquaporin 1 (AQP1) as a putative BAT silencer regulatory factor and show through the generation of the first BAT-specific aquaporin-1 knockout (AQP1-KO) mouse that BAT can be activated under environmental silencing conditions. We further show that these mice are protected against diet-induced obesity, with improved glucose tolerance, and increased energy expenditure. These findings highlight AQP1 as a promising therapeutic target in the emerging research field of BAT silencers.

## Introduction

The increasing prevalence of obesity in the United States alone has reached epidemic proportions and has now resulted in nearly 74% of US adults being classified as overweight or obese (1–3). Obesity, which increases the risk for co-morbidities such as type 2 diabetes, cardiovascular disease, stroke, and some cancers, is now one of the leading causes of preventable death in the United States (4–8). Current treatment options to combat obesity-associated metabolic dysfunction, such as dieting or bariatric surgery, can be ineffectual in the long term or have severe side effects (9, 10). Therefore, there is a critical need for novel treatments to combat this obesity epidemic.

Recent efforts have focused on the activation of brown adipose tissue (BAT) through the non-shivering thermogenesis (NST) program as a possible therapeutic option to correct the metabolic dysfunction associated with diet-induced obesity (DIO). Human BAT activation through environmental cold exposure or β3 adrenergic stimuli converts chemical energy stores into heat by uncoupling the electron transport chain in mitochondria, creating a futile cycle to increase energy expenditure (11–16). The activation of NST has resulted in significant weight loss in rats and mice in preclinical studies (17) and the presence and activation of BAT is positively correlated with improved metabolic health in humans (18–20). Most importantly, previous studies have shown that activating brown fat in obese human patients has resulted in weight loss and is inversely correlated with BMI (21–24).

However, the major limitation to leveraging brown fat as a therapeutic option to overcome metabolic dysfunction associated with obesity is that human brown fat activation is extremely transient in nature. Brown fat is silenced in humans for most of adult life under physiological temperature conditions. Though the mechanism of cold exposure-induced activation is retained in humans, constant stimulus is required to maintain NST, thus limiting its therapeutic potential. A previous study demonstrated that all metabolic benefits seen in human subjects after a 2-hour cold exposure period to activate BAT were immediately lost following a re-warming period back to ambient temperatures (25). Since then, there have been a few studies aiming to characterize regulatory factors that restrict brown fat activation, such as the identification of the cardiotrophin-like cytokine factor 1 (CLCF1) signaling pathway. This signaling pathway dampens non-shivering thermogenesis and disrupts metabolic homeostasis by inhibiting mitochondrial biogenesis in brown adipocytes, resulting in a loss of brown fat identity (26). On a larger scale, however, the mechanisms that silence brown fat in physiological ambient human temperature conditions remain poorly understood.

Defining the regulatory factors that govern brown fat silencing is a promising avenue for research. Ablation of these silencers could potentially maintain BAT in an active state, overcome the metabolic disturbances that are associated with obesity, and improve metabolic health. To this end, we have captured the first BAT silencer proteome from wild-type (WT) mice exposed to environmental thermoneutral (BAT silencing) conditions compared to cold (BAT activating) conditions. Here, we discovered the protein aquaporin-1 (AQP1), a canonical water channel protein that harbors thermogenic silencing regulatory potential. AQP1 is one of 13 members of the aquaporin family that functions predominantly as a membrane-bound transport protein for water and small ions (27–31). Specific members of the aquaporin family have been shown to be critical players in obesity and adipose tissue metabolism (32–37). For instance, aquaporin-7 (AQP7) is an aquaglyceroporin that is permeable to both water and glycerol and thus facilitates glycerol efflux from adipocytes. AQP7 dysregulation has been positively correlated with obesity onset and is associated with adipocyte differentiation, insulin response and fat metabolism (32, 38, 39). However, AQP1, the archetype of water channels, is currently unannotated for a functional role in BAT and obesity-associated metabolic dysfunction.

Through the generation of the first BAT-specific AQP1 knockout mouse, we demonstrate that AQP1 expression is not only inversely correlated to UCP1 expression, but also highly expressed under brown fat silencing conditions. We further show that ablation of AQP1 in BAT confers robust protection against DIO and normalizes blood glucose levels by upregulating the electron transport chain (ETC) and mitochondrial translation program under thermoneutral BAT silencing conditions. Our work demonstrates that ablation of AQP1 in BAT promotes the activation of BAT under ambient temperature conditions. This suggests that AQP1 plays a key role in the regulation of brown fat silencing and may be an effective translational therapeutic target to combat obesity-driven metabolic dysfunction.

## Results

### AQP1 expression is significantly upregulated under BAT silencing environmental conditions

To capture thermogenic brown fat silencer candidates, we speculated that a silencer regulatory factor would likely be most abundant and active in the thermoneutral (TN) environmental state when brown adipose tissue (BAT) is silenced compared to cold environmental conditions when brown fat is active (Fig. 1A). We therefore analyzed the proteome in BAT from 6-week-old mice exposed to either cold (CD) or TN temperature conditions for 7 days to capture proteins significantly elevated in TN conditions but low in CD. We then collected BAT from these mice and performed tandem mass tagged (TMT) proteomic profiling. Our proteomic screen revealed 137 proteins that were significantly upregulated in TN conditions relative to CD (Fig. 1B). We subsequently filtered these candidates by statistical significance, orthologous expression in human BAT for future translational potential, and novelty (40–42). This revealed five protein silencer candidates, with Aquaporin-1 (AQP1) being the most significant (Fig. 1C-D). Subsequent protein profiling validation of our proteomics findings confirmed that indeed, AQP1 and UCP1 were observed to have inverse protein expression dynamics in CD versus TN conditions (Fig. 1E-G), thus confirming AQP1 as our top candidate to further investigate potential BAT silencing mechanisms.

**Fig 1.**
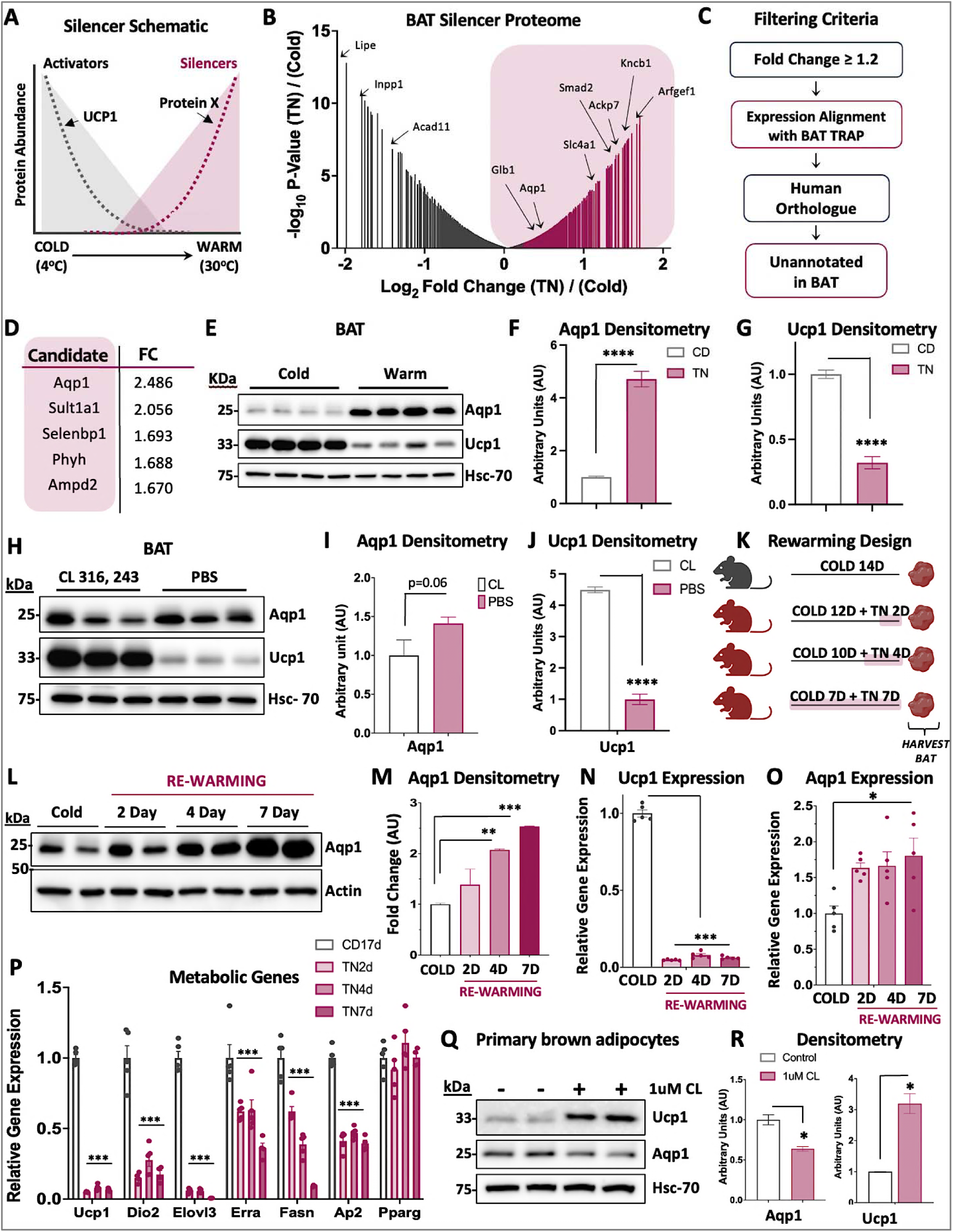
Aquaporin 1 levels increase upon NST silencing. (A) Schematic of proteomic screen expected protein abundance dynamics for thermogenic activators and silencers in cold versus warm conditions in whole brown adipose tissue (BAT) isolated from 6-week-old wild-type (WT) C57BL/6J male mice. (B) Line volcano plot of WT BAT proteomics screen results highlighting putative thermogenic activators and silencers (TN7d vs CD7d, n=3/ group). (C) Simplified diagram of filtering criteria used to identify possible thermogenic silencer candidates. (D) Table displaying the protein names of the top five putative thermogenic silencer candidates along with their corresponding fold change in warm vs cold conditions. (E) Representative immunoblot of BAT isolated from 6-week-old WT male mice exposed to cold (CD) or thermoneutral (TN) for 7 days (n=4) to repeat and validate proteomics screen results. (F) Densitometry quantification of AQP1 protein expression normalized to loading control HSC70 (Heat shock cognate 71 kDa protein) from 1E immunoblot in CD and TN 7d conditions. (G) Densitometry quantification of UCP1 protein expression normalized to loading control HSC70 from 1E immunoblot in CD and TN 7d conditions. (H) Representative immunoblot of BAT from 6-week-old male WT mice treated with PBS vehicle or 1mg/kg CL 316,243 (CL) injection for 14 days (n=5). (I) Densitometry quantification of AQP1 protein expression normalized to loading control HSC70 from 1H immunoblot under PBS or CL treated conditions. (J) Densitometry quantification of UCP1 protein expression normalized to loading control HSC70 from 1H immunoblot under PBS or CL treated conditions. (K) Illustration of BAT harvested from WT male mice to test dynamics during re-warming period: Cold 14d, Cold 12d + TN 2d, Cold 10d + TN 4d, and CD 7d + TN7d. (L) Representative immunoblot of BAT isolated from WT male mice exposed to CD 7d, compared to mice exposed to CD for at least 7 days followed by a re-warming period of 2, 4, or 7 days in TN (n=5) to repeat and validate proteomics screen results. (M) Densitometry quantification of AQP1 protein expression normalized to actin from Fig. 1F immunoblot. (N) UCP1 relative mRNA expression from proteomics validation experiment. (O) AQP1 relative mRNA expression from proteomics validation experiment. (P) Relative mRNA expression of thermogenic biomarkers and related metabolic genes from proteomics validation experiment (n=5). (Q) Representative immunoblot of primary brown adipocytes isolated from 3-week-old WT male mice and were differentiated for 7 days and treated with PBS vehicle or 1uM CL for 24h (n=2 technical duplicates). (R) Densitometry quantification of AQP1 and UCP1 protein expression normalized to HSC70 loading control from 1Q immunoblot. Densitometry and qPCR data represented as mean + SEM. Significance is denoted as *p<0.05, **p<0.01, ***p<0.001, ****p<0.0001 by Student’s t test.

To observe whether this inverse temperature-dependent expression dynamic between UCP1 and AQP1 was mediated by β3 adrenergic receptor signaling, we injected mice with either PBS or CL 316,243 (CL), a β3 adrenoceptor agonist to activate BAT, for 14 days. We observed a similar inverse protein expression dynamic between AQP1 and UCP1 with CL treatment, although the increase in AQP1 protein levels under pharmacological BAT silencing was not as robust as was observed with TN exposure. This suggests that the regulation of AQP1 under BAT silencing conditions is not solely through β3 adrenergic receptor signaling (Fig. 1H-J).

To more clearly define the AQP1 protein expression dynamics over time, we exposed mice to CD conditions for a minimum of 7 days for maximal BAT activation and then re-exposed them to thermoneutrality for 2, 4, and 7 days to represent acute, moderate, and chronic brown fat silencing conditions (Fig. 1K). As observed with our proteomics results, AQP1 protein levels were low in cold and gradually increased at the protein level upon BAT silencing (Fig. 1L and 1M). This expression dynamic was unique to BAT as there was no change in AQP1 expression in beige inguinal white adipose tissue (iWAT) or kidney—an abundant source of AQP1 (*SI Appendix*, Fig. S1A-B). Furthermore, we found that while *Ucp1* and other classical thermogenic and mitochondrial markers were silenced when reintroduced to TN conditions, *Aqp1* had an inverse gene expression pattern that increased upon BAT silencing (Fig. 1N-P and *SI Appendix*, S1C). Interestingly, other aquaporins, such as *Aqp7* and *Aqp11*, had significant upregulated gene expression in CD, but lower gene expression in TN conditions, which is the opposite gene expression dynamic of AQP1 (Fig. S1D).

In order to determine if AQP1 exhibited the same dynamic pattern through adrenergic signaling, we modeled chemical silencing conditions by first injecting CL for 7 days and then replacing this treatment with PBS administration for 2, 4, and 7 days to model the same acute, moderate, and chronic silencing conditions as was done with cold exposure (Fig. 1K). We then profiled AQP1 in BAT at the protein and mRNA level (*SI Appendix*, Fig. S1E-H) and found similar, but trending, dynamics to what we observed under CD-TN rewarming conditions, which suggests that β3 adrenergic signaling partially contributes to the regulation of AQP1 expression patterns.

BAT is an extremely heterogenous tissue with many cell types and AQP1 is also expressed in other cell types other than adipocytes, such as endothelial cells and macrophages (43–47) (*SI Appendix*, Fig. S2A). To examine whether the brown adipocyte population was driving the AQP1 dynamics observed at the tissue level, we cultured and differentiated primary murine brown adipocytes from male WT mice and stimulated them with CL or PBS control in culture to simulate pharmacological NST active and silencing conditions in culture respectively. We show at both the protein and mRNA level that AQP1 expression was high under PBS conditions (NST silenced) and significantly lower upon CL administration (NST activated). Consistent with previous observations at the tissue level, the expected UCP1 dynamic was inverse to AQP1 expression, suggesting that the enhanced AQP1 expression pattern and dynamic in BAT silencing conditions is driven by brown adipocytes specifically (Fig. 1Q-R and *SI Appendix*, S1I-J). Thus, AQP1 is robustly upregulated in BAT silencing conditions but significantly downregulated in BAT active conditions and is inversely correlated to BAT activators, which matches the expected expression pattern of a putative BAT silencer.

### BAT-specific AQP1-KO mice are protected against diet-induced obesity and metabolic dysfunction *in vivo*

We created the first BAT-specific AQP1 knockout (AQP1-KO) mouse model by crossing homozygous AQP1-Flox mice with mice expressing a hemizygous UCP1 promoter Cre recombinase (Fig. 2A-B). Given the wide range of AQP1 expression across different cell types in BAT, we predicted and confirmed that the knockout of AQP1 was masked at the whole tissue level but was validated in primary AQP1-Flox and KO brown adipocyte cultures (43, 44) (Fig. 2C-D and *SI Appendix*, S2A).

**Fig 2.**
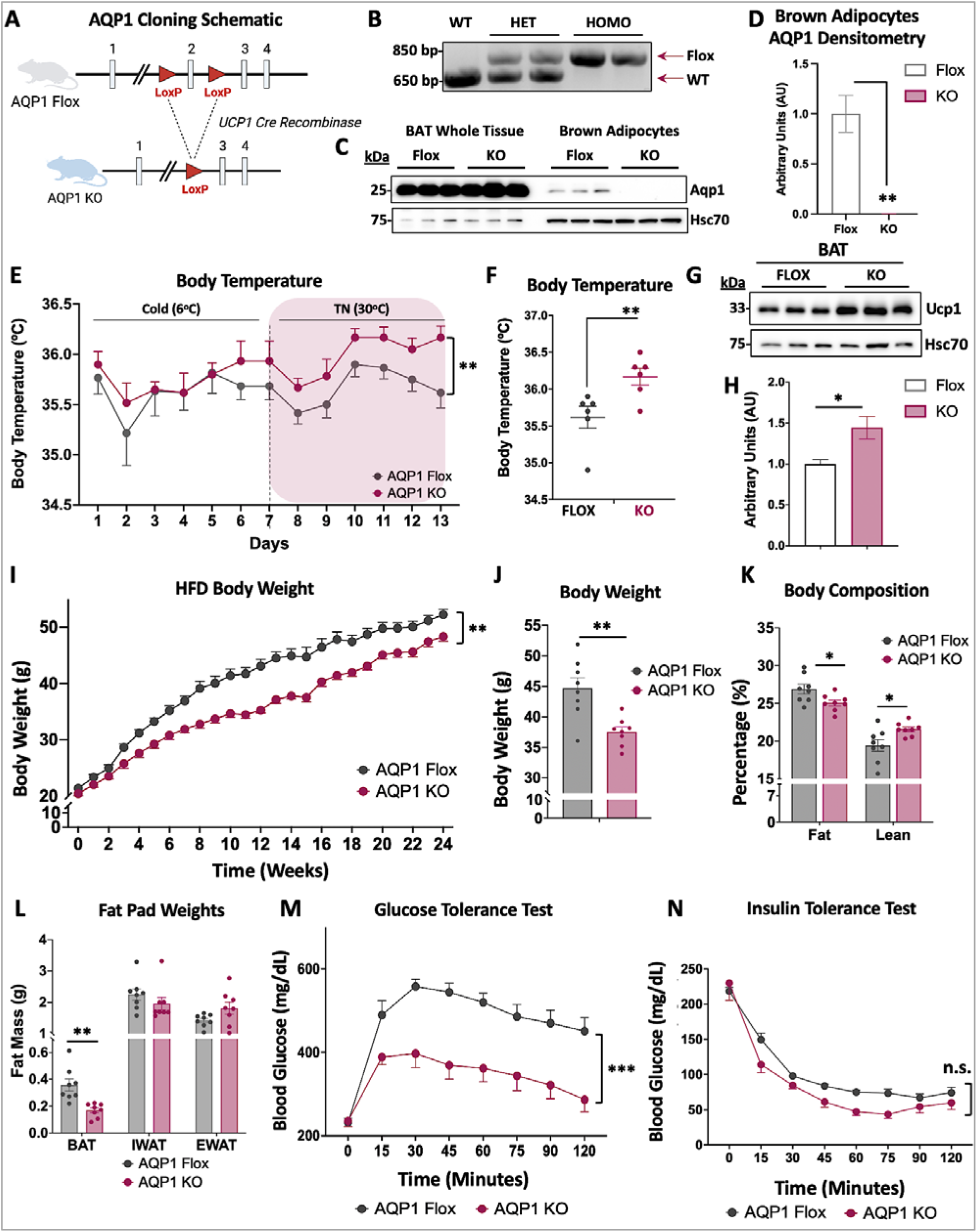
BAT-specific AQP1-KO protects against diet-induced obesity. (A) Schematic of BAT-specific AQP1-KO (AQP1-KO) mouse model breeding strategy. (B) Example of genotyping of WT, heterozygous or homozygous AQP1-Flox (AQP1-Flox) mice. (C) Representative immunoblot of AQP1 expression in AQP1-Flox and AQP1-KO in whole BAT versus differentiated primary brown adipocytes from 3-week-old AQP1-Flox male mice treated with adenovirus GFP or Cre. (D) Densitometry quantification of AQP1 protein expression normalized to HSC70 from 2C immunoblot. (E) Body temperature measured over time and taken by daily rectal body temperature measurements through the CD 7d followed by TN 7d experiment (n=6). (F) Quantification of rectal body temperature of AQP1-Flox and KO mice on TN 7d from 2E. (G) Representative immunoblot of UCP1 expression in BAT from AQP1-Flox and AQP1-KO mice. (H) Densitometry quantification of UCP1 expression normalized to HSC70 from 2G. (I) Body weight of AQP1-Flox and AQP1-KO male mice measured weekly over 24 weeks of high-fat diet (HFD) housed at 30°C (n=8). (J) Body weight of AQP1-Flox and KO mice measured at week 15 of HFD (n=8). (K) Body composition measured by nuclear magnetic resonance (NMR) to measure fat and lean percentage (n=8). (L) Fat pad weights from brown adipose tissue (BAT), inguinal white adipose tissue (iWAT), and epididymal white adipose tissue (eWAT) measured after 25 weeks of HFD. (M) Glucose tolerance test (GTT) conducted on week 17 of HFD for AQP1-Flox and KO mice. Mice fasted for 16h and injected with 1.5mg/kg glucose. Measurements were taken by glucometer in duplicate every 15min over a 120 min period (n=8). (N) Insulin tolerance test (ITT) conducted on week 20 of HFD for AQP1-Flox and KO mice. Mice fasted for 4h and injected with 1U/kg insulin. Measurements were taken by glucometer in duplicate every 15min over a 120 min period (n=8). Endpoints of body weight, GTT, ITT time courses and all bar graphs are represented as mean + SEM. Significance is denoted as *p<0.05, **p<0.01, ***p<0.001 by Student’s t test.

We then hypothesized that ablation of AQP1 in BAT would activate NST under ambient temperature conditions when BAT is normally silenced. To investigate this, we exposed chow-fed AQP1-Flox and AQP1-KO mice to CD challenge for 7 days followed by exposure to TN for 7 days, taking body temperature measurements daily. There were no differences in body temperature between the two groups in CD, but interestingly, we observed that AQP1-KO mice had significantly higher body temperature and UCP1 protein levels in their BAT tissue under TN conditions, suggesting that ablation of AQP1 allowed NST to remain active under BAT silencing environmental conditions (Fig. 2E-H). Curiously, there were no significant differences in UCP1 or other NST or mitochondrial biomarker expression at the mRNA level, suggesting possible post transcriptional regulation to keep NST active (*SI Appendix*, Fig. S2B-D).

Given that the chow-fed AQP1-KO seemed to have activated BAT in TN conditions, we speculated that ablation of AQP1, as a putative NST silencer in BAT, should also confer protection against DIO under ambient TN temperature conditions when BAT is silenced. We therefore placed 6-week-old AQP1-Flox and AQP1-KO mice on 60 kcal% high fat diet for 24 weeks in TN conditions (30°C) to prevent activation of NST. Throughout the 24-week HFD feeding regimen, AQP1-KO mice gained significantly less body weight gain compared to their AQP1-Flox control group counterparts (Fig. 2I-J). This protection occurred with no differences between the two groups in food or water consumption (*SI Appendix*, Fig. S2E-F). At the body composition level, the lower body weight in the AQP1-KO stem from significant reductions in fat mass with a corresponding increase in lean mass (Fig. 2K) but no changes in free fluid (*SI Appendix*, Fig. S2G). The decrease in fat mass at the whole-body level in the AQP1-KO mice was likely driven by the decrease in BAT tissue mass, as there were no significant differences in inguinal or epidydimal white adipose tissue masses (iWAT and eWAT respectively) (Fig. 2L).

To determine if the protection against HFD feeding in the AQP1-KO mice also confers improved glucose homeostasis, we performed glucose and insulin tolerance assessments. Impressively, we found that AQP1-KO mice had significant improvements in glucose tolerance relative to control mice (Fig. 2M). Yet, there were no differences in insulin sensitivity between the control and AQP1-KO mice (Fig. 2N) or plasma leptin concentration (*SI Appendix*, Fig. S2H). This would suggest that the enhanced glucose tolerance of the AQP1-KO mice under HFD conditions is not being driven by enhanced insulin sensitivity or alterations in hormonal leptin levels. Taken together, we show that ablation of AQP1 in BAT protects mice against DIO by reducing fat mass, increasing lean mass and significantly improving glucose homeostasis. Furthermore, improvements in metabolic parameters were achieved under environmental conditions where BAT is silenced, suggesting that AQP1 may play a role in the regulation of BAT silencing.

### AQP1-KO increases whole body energy expenditure

Following 24 weeks of HFD feeding, AQP1-Flox and KO mice were placed in Promethion metabolic cages to evaluate their whole-body metabolism. We found that the AQP1-KO mice had significantly increased energy expenditure compared to the control mice, which was likely a key factor in their protection against metabolic dysfunction (Fig. 3A-D). The increase in energy expenditure occurred with no changes in respiratory exchange ratio, total food or water consumption, or locomotive movement, suggesting that this increase in energy expenditure was driven by BAT activation (Fig. 3E and *SI Appendix*, Fig. S3A-F). Curiously, there were no significant differences in UCP1 or mitochondrial biomarkers at the mRNA or protein level, suggesting that the increase in energy expenditure under HFD conditions may be driven by increases in UCP1 activity rather than absolute protein levels (Fig. 3F and *SI Appendix*, Fig. S3G-H). As the AQP1-KO is specifically in the brown adipocyte cell population, we then wanted to understand whether AQP1 ablation in this population was driving the increased energy expenditure and protection against obesity phenotype in a cell-autonomous manner. We therefore cultured and differentiated primary AQP1-Flox and AQP1-KO brown adipocytes to determine the impact of AQP1 ablation *in vitro*. Consistent to what was observed at the whole tissue level, there were no significant differences in either UCP1 or core mitochondrial proteins in the oxidative phosphorylation (OXPHOS) pathway (Fig. 3G-H). We then evaluated whether the AQP1-KO brown adipocytes contributed directly to the improved glucose homeostasis observed in the HFD mice by measuring cellular glucose uptake rate. We discovered that the AQP1-KO primary brown adipocytes indeed had a significantly higher rate of glucose uptake compared to the AQP1-Flox cells (Fig. 3I). This suggests that brown adipocytes that lack AQP1 can clear more glucose from circulation, which aligns with the enhanced glucose blood clearance that was seen *in vivo* (Fig. 2L).

**Fig 3.**
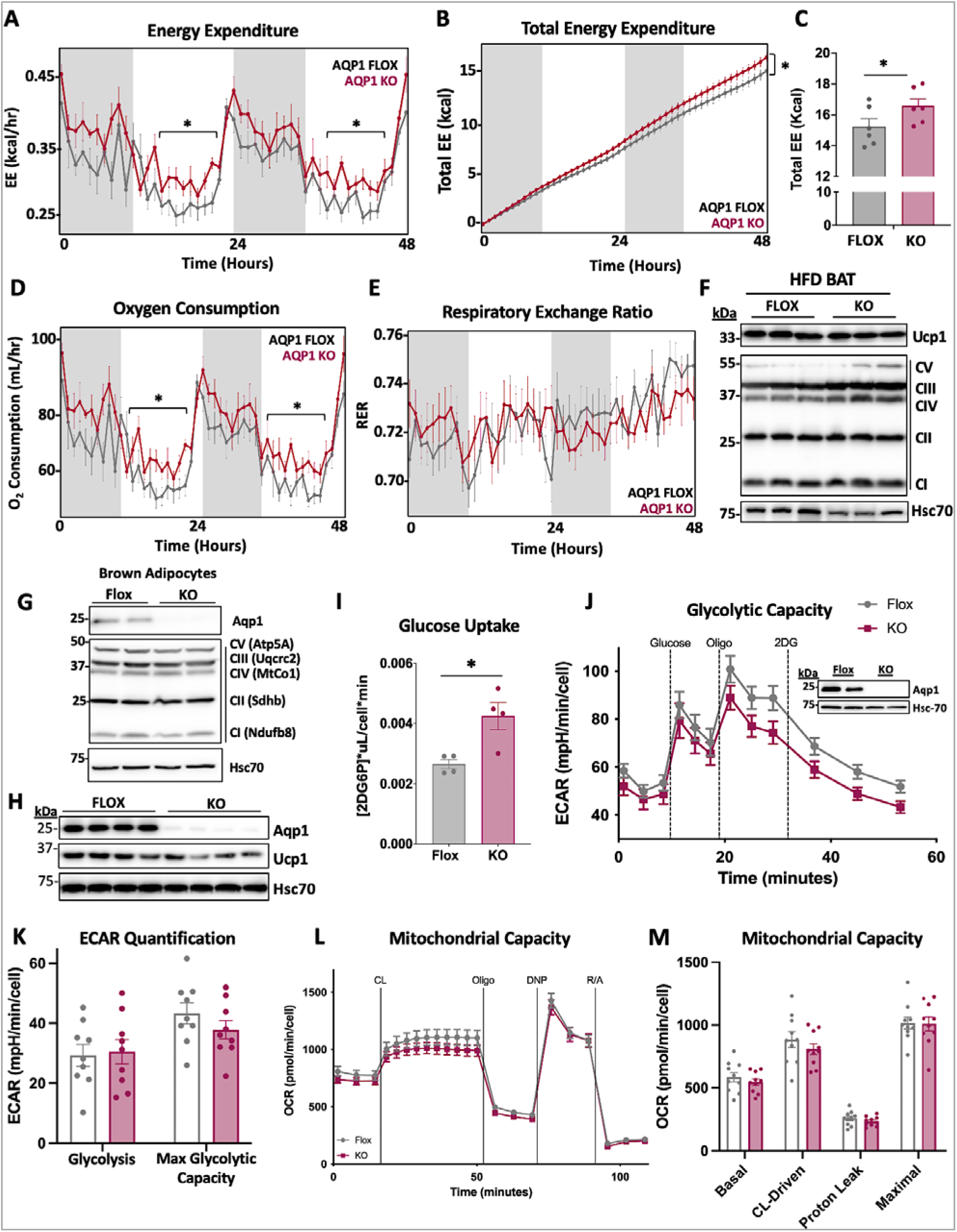
AQP1-KO increases whole body energy expenditure but not by traditional UCP1-dependent NST. (A-E) Metabolic data obtained from Promethion metabolic cages of AQP1-Flox and AQP1-KO male mice single-housed at 30°C for a 48h period after 24 weeks of HFD (n=6). (A) Energy expenditure. (B) Total energy expenditure. (C) Bar graph displaying total energy expenditure of AQP1-Flox and AQP1-KO at 48h endpoints. (D) Oxygen consumption. (E) Respiratory exchange ratio (RER). (F) Representative immunoblot of UCP1 and total OXPHOS expression in BAT harvested from AQP1-Flox and AQP1-KO male mice after 25 weeks of HFD. (G-L) *In vitro* data obtained from cultured primary brown adipocytes isolated from 3-week-old AQP1-Flox male mice, treated with adenovirus GFP or Cre for 72h, and differentiated for 5-7 days. (G) Representative immunoblot of differentiated primary brown adipocytes validating AQP1-KO in cell culture and displaying total OXPHOS expression (n=2). (H) Representative immunoblot of differentiated primary brown adipocytes validating AQP1-KO in cell culture and displaying UCP1 expression (n=4). (I) Rate of glucose uptake bar graph measured from differentiated primary brown adipocytes (n=4). (J) Glycolytic capacity with the Seahorse Bioanalyzer in the AQP1-Flox and AQP1-KO primary brown adipocytes (n=10). Representative immunoblot showing corresponding validation of AQP1-KO in samples. (K) Quantification of glycolytic capacity of AQP1-Flox and AQP1-KO primary brown adipocytes (n=9). (L) Mitochondrial respiratory capacity of AQP1-Flox and AQP1-KO primary brown adipocytes (n=10). (M) Quantification of mitochondrial respiratory capacity of AQP1-Flox and AQP1-KO primary brown adipocytes (n=10). (A-E) was analyzed by ANCOVA; All other Figures unless otherwise indicated are data represented as mean ± SEM. *p < 0.05 by Student’s t test.

We postulated that the increased glucose uptake rate could be used as a substrate for increased cellular bioenergetic capacity. We therefore measured cellular glycolytic and oxidative mitochondrial capacities in the AQP1-Flox and KO brown adipocytes but surprisingly we found no differences between the KO and the control group (Fig. 3J-M). This indicated that the increased glucose uptake was not supporting an increase in glycolytic or oxidative capacity in the brown adipocyte population. Thus, the increase in whole body energy expenditure observed in Fig. 3A is not solely driven by the brown adipocyte population. This implies that another mechanism, such as crosstalk with other cell types or among tissues at the whole-body level, may be required to drive this *in vivo* phenotype.

### AQP1-KO mice have upregulated mitochondrial metabolism genes on HFD

Ablation of AQP1 in brown adipocytes increases NST under environmental conditions where BAT is normally silenced *in vivo,* but this effect was not observed in *ex vivo* cultured brown adipocytes in a cell-autonomous manner. To mechanistically define how ablation of AQP1 in BAT protects against metabolic dysfunction *in vivo*, we interrogated the BAT proteome from HFD AQP1-KO and control AQP1-Flox mice and performed shotgun proteomic analysis (Fig. 4A). Gene ontology analysis from the top differentially expressed proteins revealed significant upregulation in the oxidative phosphorylation (OXPHOS) and mitochondrial translation pathways in the BAT depots (Fig. 4B). Indeed, a host of core electron transport chain proteins and mitochondrial ribosomes were upregulated in the BAT of AQP1-KO mice compared to Flox control mice, which also showed significant increases at the transcript level (Fig. 4C and D). Taken together, we determined that ablation of the putative NST silencer AQP1 in BAT significantly upregulates mitochondrial energy metabolism and translation at the BAT tissue level, which likely contributes to increased energy expenditure, protection against diet-induced obesity, and normalized blood glucose profiles *in vivo*.

**Fig 4.**
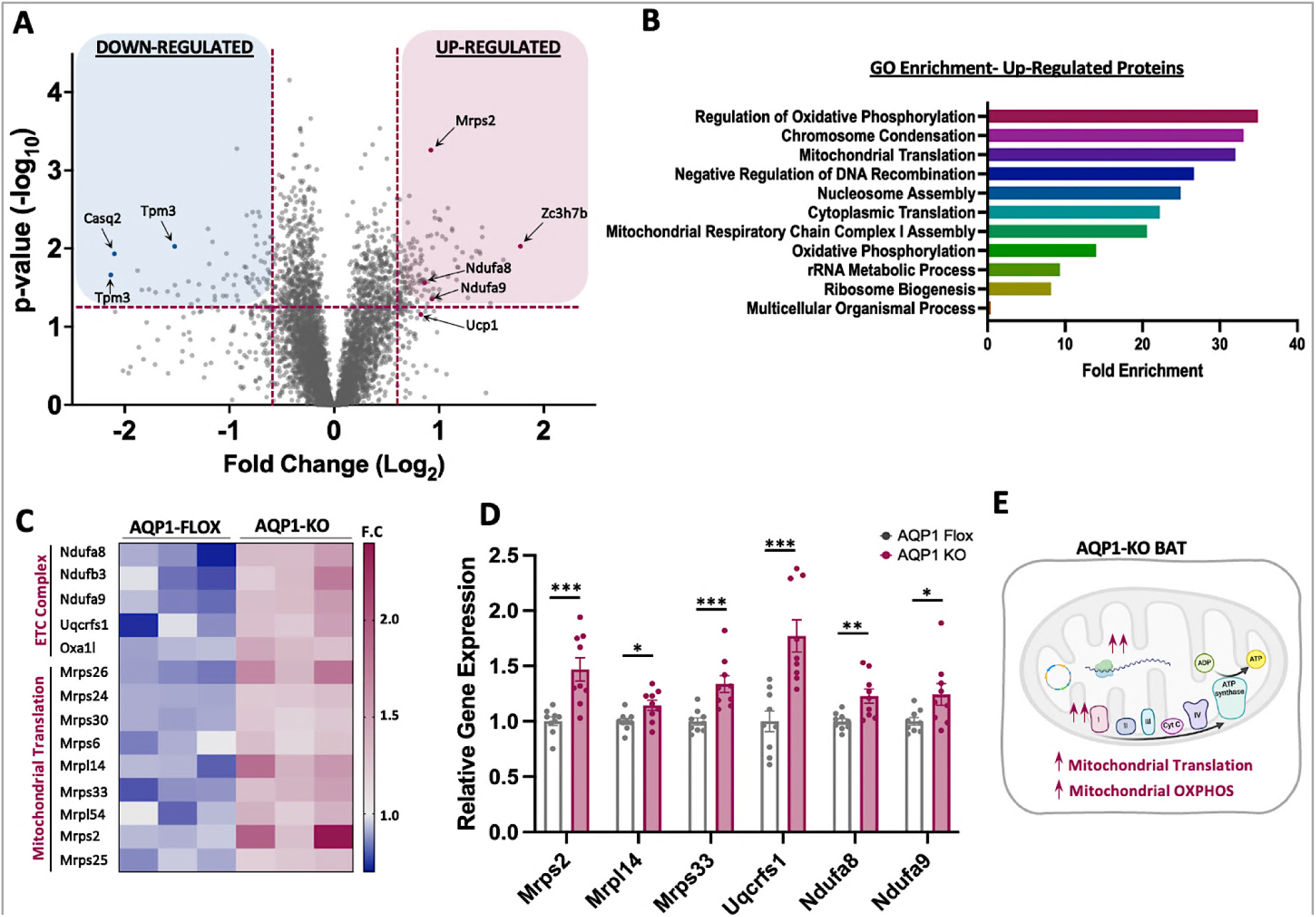
AQP1 ablation on HFD enhances mitochondrial metabolism. (A) Volcano plot of BAT harvested from AQP1-Flox and AQP1-KO male mice after 25 weeks of HFD. Proteomic results highlight significantl upregulated and downregulated proteins in AQP1-KO mice versus AQP1-Flox mice (n=4/ group). (B) Gene Ontology (GO) analysis of top ten upregulated pathways based on the top 197 significantly upregulated proteins from 4A proteomic screen. (C) Heat map displaying specific upregulated electron transport chain (ETC) complex and mitochondrial translation proteins from 4A proteomic screen and 4B GO analysis. (D) Relative mRNA expression of three mitochondrial translation and three ETC complex proteins. (E) Schematic of proposed mechanism of how AQP1 ablation affects AQP1-KO mice relative to AQP1-Flox mice. Relative mRNA expression data represented as mean + SEM. Significance is denoted as *p<0.05, **p<0.01, ***p<0.001 by Student’s t test.

## Discussion

It is well known that the activation of BAT is extremely temporal in nature and rapidly becomes silenced once the activation stimulus is removed (25, 48–56). This phenomenon continues to be a challenge in the field as many groups continue to find novel methods to sustain the activation of BAT (57). Despite this challenge however, very little is known about the regulation of BAT silencing. Here, we discovered the water channel protein AQP1 as a putative BAT silencing regulatory factor from a proteomic screen conducted under brown fat environmental silencing conditions. We show that ablation of AQP1 robustly protects against DIO and metabolic dysfunction in mice by increasing whole-body energy expenditure through an upregulation of the oxidative phosphorylation and translation proteins in brown fat.

Select aquaporins have well-documented roles in metabolism. The most well-studied aquaporin in this realm is the aquaglyceroporin AQP7, which transports both water and glycerol and thus facilitates glycerol efflux from adipocytes. AQP7 dysregulation has been tied to the onset of obesity in mice due to impaired glycerol dynamics, triglyceride accumulation, and metabolic dysfunction (32, 35, 38, 39, 58, 59). While AQP1 has been previously studied in other contexts such as kidney disease and cancer (30, 60, 61), little is known about AQP1 in adipose tissue specifically. A recent study by Costa *et al*. discovered that fasting conditions increased AQP1 expression and increased net ultrafiltration across the peritoneal membrane, which indicates that AQP1 may be regulated by nutrient status (59). Furthermore, Madonna *et al*. demonstrated that high glucose and high mannitol increased AQP1 expression in human aortic endothelial cells and impaired insulin signaling, but AQP1 silencing reversed this effect (62). Thus, these studies strongly suggest that AQP1 may play a role in regulating metabolism in response to nutrient status and to date, we are the first to report a such a relationship between AQP1 and energy metabolism in brown fat.

AQP1 expression, at both the transcript and protein level, is highly upregulated under BAT silencing environmental conditions and is inverse to the expression of classical thermogenic proteins such as UCP1. This inverse expression pattern to classical BAT activators is unique to BAT as other tissues, such as kidney, that have high levels of AQP1 do not display this pattern (Fig. S1B). Motivated by our findings in WT mice, we generated the first BAT-specific AQP1-KO mouse model by crossing AQP1-Flox mice with mice expressing hemizygous UCP1 Cre recombinase. We hypothesized that AQP1 ablation as a putative silencer regulatory factor may alleviate BAT silencing mechanisms and promote the activation of NST in TN conditions, thereby protecting mice against DIO. Impressively, we observed an increase in body temperature *in vivo* under TN conditions in chow-fed mice and significant decreases in diet-induced weight gain and improved glucose tolerance under HFD conditions in the AQP1-KO mice, supporting the possible role of AQP1 as a BAT silencer regulatory factor. Curiously, despite the drastic improvements in glucose clearance, we found no difference in insulin tolerance. One interesting explanation for this finding could be that AQP1 ablation mitigates insulin resistance by blocking transmembrane transport of hydrogen peroxide (H_2_O_2_). Previous literature has reported that H_2_O_2_ overproduction from myocellular mitochondria under DIO conditions has been linked to impaired insulin signaling and glucose transport regulation (63–66). Most recently, Montiel *et al*. found that AQP1 can facilitate the transmembrane passage of H_2_O_2_ through its water channel, revealing its new function as a “bona fide peroxiporin” (67). If DIO causes the overproduction of H_2_O_2_, then the absence of AQP1 could limit H_2_O_2_ transport in mice on HFD. Thus, it is possible the HFD-fed AQP1-KO mice regained insulin sensitivity since they do not have highly reactive hydroxyl radicals that could significantly diminish IRS/PI3K/Akt-dependent insulin signaling.

AQP1-KO mice on HFD had increased energy expenditure and oxygen consumption that is directly correlated with an upregulation in ETC complex and mitochondrial translation protein expression. At the *in vitro* level, we also found that AQP1-KO primary brown adipocytes had an increased rate of glucose uptake. These findings suggest that NST may be contributing to the protection against DIO that we observed *in vivo*. However, this increase in energy expenditure does not seem to be driven by traditional UCP1-dependent NST as there were no differences in UCP1 or total OXPHOS protein expression between AQP1-Flox and KO, both at whole BAT and primary brown adipocyte levels. While it is possible that protein activity, but not expression, was enhanced due to an increase in substrate utilization, we cannot rule out the possibility that UCP1-independent mechanisms, such as enhanced creatine or succinate futile cycling, could also be driving an increase in energy expenditure.

Surprisingly, when we analyzed cellular respirometry in AQP1-KO primary adipocytes compared to Flox controls, there were also no observable differences in glycolytic or mitochondrial oxidative capacity in AQP1-KO primary brown adipocytes. This suggests that the increase in energy expenditure *in vivo* is retained by the brown adipocyte in culture and may not be a cell-autonomous phenomenon. One possible explanation for this finding is the inherent heterogeneity of cell types in BAT. Sun *et al*. previously published single nuclei RNA sequencing (snRNAseq) data that revealed cell-type heterogeneity in BAT (43). Thus, if cell-to-cell communication between brown adipocytes and other cell types (e.g. endothelial cells, immune cells, vascular cells, neuronal cell) (68) is required to observe this phenomenon, this could explain why we observed no differences in NST biomarker expression or functional respiration in isolated cultured brown adipocytes.

Another possible explanation for this phenomenon, beyond NST activation, is that the AQP1 protein itself is transporting oxygen. Previous studies have found that the AQP1 protein can facilitate oxygen transport (69, 70), but AQP1 function, especially in this context, has yet to be annotated in BAT. It is possible that in whole BAT, AQP1 ablation in brown adipocytes may be upregulating the AQP1 expression in other cell types to sustain increased oxygen transport to support thermogenesis. Therefore, because there are no other cell types present to compensate for the loss of AQP1 expression *in vitro*, this may be why we did not observe any differences in functional respiration at the cellular level. A third possibility is that AQP1’s function as a water channel may directly influence cellular stiffness or adipocyte shape (71), which has been shown to impact brown adipocytes’ ability to perform uncoupled respiration. However, further investigation into AQP1 gas permeability, water channel and ion channel function are required to understand the full mechanism of how AQP1 ablation in BAT protects mice against DIO.

Taken together, these data would suggest that AQP1 ablation in BAT can protect AQP1-KO mice against DIO by increasing energy expenditure through upregulation of ETC complexes and mitochondrial translation gene expression. Targeting putative silencers of BAT activation to sustain NST metabolic benefits in ambient temperatures proposes a noninvasive, effective molecular therapeutic strategy to combat obesity-associated metabolic dysfunction. Thus, the findings of this paper represent the first documented phenotypes of BAT-specific AQP1 ablation and determined that AQP1 is a promising therapeutic target for inhibition to protect against DIO and metabolic disease.

## Materials and Methods

### Mouse Studies

All animal experiments were approved by the Institutional Animal Care and Use Committee (IACUC) at Cornell University. To generate the BAT-specific AQP1-KO transgenic mice, homozygous floxed AQP1^Flox/Flox^ mice were obtained from Dr. Olivier Devuyst from the University of Zurich (29, 31, 72). These mice were crossed with hemizygous UCP1^Cre/WT^ mice (Jackson #024670) to generate the final study genotypes AQP1^Flox/Flox^ and AQP1^Flox/Flox^/UCP1^Cre/WT^ representing the AQP1-Flox control and AQP1-KO mice respectively. For all animal studies, 6-week-old male and female mice (female data not shown) were used. The specific number of mice used is indicated in each Fig.. All mice were maintained on 14h light and 10h dark cycles and fed *ad libitum* with either standard irradiated rodent chow or 60 kcal% high fat diet (HFD) (Research Diets D12492). HFD treatment was initiated in mice starting at 6 weeks of age. These mice were housed at thermoneutral (TN) temperature (30°C) for 8-24 weeks with weekly weight measurements and cage changes. For temperature manipulation experiments, mice were born and reared at room temperature (RT-22°C) until 6 weeks of age. Mice were then acclimated at thermoneutrality (TN-30°C) for 2-3 days, followed by exposure to room temperature (RT-22°C) for 1 day, before cold (CD) exposure (6.5°C) for 7 days, and then returned to TN for 7 days. Rodent rectal body temperature was measured daily using a Thermalert temperature probe (Physitemp) coated in 100% glycerol. For reverse thermogenesis experiments (CD followed by TN), 6-week-old wild-type (WT) C57BL/6J (Jackson #000664) and AQP1 transgenic mice were single-housed in rodent incubators and exposed to a constant temperature of 6.5°C followed by 30°C for indicated timepoints. For CL 316,243 injections, 6-week-old WT mice were injected with PBS vehicle or CL 316,243 at 1mg/kg daily for 14 days in TN.

### Murine Primary Brown Adipocyte Cell Culture

Primary brown adipocytes were isolated from brown adipose tissue (BAT) in 3-week-old male wild-type (WT) C57BL/6J mice or AQP1 transgenic male mice. BAT pads from 5 mice were pooled after dissection, minced thoroughly for 5 min, and digested in 15 ml of BAT dissociation buffer (123mM NaCl, 5mM KCl, 1.3mM CaCl2, 5.0 mM Glucose, 100 mM HEPES, 4% BSA and 1.5 mg/ml collagenase B) for 30min at 37°C with constant shaking. The cell suspension was filtered with a 100 µm cell strainer and centrifuged at 600 g for 5min. The pellet was then suspended in adipocyte culture medium (DMEM/ F12 with 10% FBS, 1% PenStrep), filtered with a 40 µm cell strainer, centrifuged at 600 g for 5min, and resuspended in culture medium and plated in 2% gelatin coated 10 cm polystyrene cell culture dishes. Preadipocytes were seeded to post confluency and differentiated with DMEM/F12 supplemented with 5µg/mL Insulin, 1µM Rosiglitazone, 1µM Dexamethasone, 0.5mM Isobutylmethylxanthine (IBMX) and 1nM T3. Cells were maintained in differentiation media for 48 h before being switched into maintenance media (5µg/mL Insulin and 1µM Rosiglitazone and 1nM T3) for 5-7 days. For primary brown adipocyte adenoviral-GFP or Cre treatment, primary AQP1-Flox brown adipocytes were treated with 1000 Multiplicity of infection (MOI) of either Adeno GFP or Adeno Cre virus for 72 h. Cells were then trypsinized and reseeded into desired assay plates and exposed overnight to another dose of adenovirus treatment at an MOI of 1000. Cells were then differentiated the next day as described above. On day 5-7 of differentiation, primary brown adipocytes were treated with PBS vehicle or 1µM CL for 24 h.

### Glucose and Insulin Tolerance Tests

The glucose tolerance test (GTT) was performed at week 17 of HFD treatment. Mice were fasted for 16 h before receiving an intraperitoneal (IP) injection of 1.5mg/kg glucose. Blood glucose levels were measured using a glucometer every 15min for a duration of 2 h after glucose administration. The insulin tolerance test (ITT) was performed on week 20 of HFD treatment. Mice were fasted for 6 h before being IP injected with 1U/kg of insulin. Blood glucose levels were measured using a glucometer every 15min for a duration of 2 h post-injection.

### *In Vivo* Indirect Calorimetry by Promethion Metabolic Cages

After 25 weeks of HFD treatment, 31-week-old AQP1 male transgenic mice, were single-housed in rodent Promethion metabolic cages (Sable Systems International) situated in a temperature-controlled cabinet set to 30°C for 7 days. Comprehensive real-time metabolic measurements such as energy expenditure (kcal/hr), oxygen consumption (VO_2_), carbon dioxide expiration (VCO_2_), locomotive movement (measured by X-Y infrared beam breaks), and respiratory exchange ratio (RER) were measured and recorded every 3 min using the Sable System data acquisition software (Promethion Live v.23.0.4). Raw data was then processed using the Sable System Macro Interpreter software (v23.6.0) and One-Click Macro systems (v2.53.2). Data was further processed using the CalR software (v1.3) (73).

### Blood Plasma Collection and ELISA Blood Leptin Determination

Plasma Analysis Mice were euthanized at 3.5L/min of CO2 for 3 min. Blood was collected through a subcutaneous intracardial puncture with a 26-gauge needle. A range of 300 to 800 microliters of blood was collected per subject and placed in Sarstedt k3 EDTA Plasma microcentrifuge tubes. Microcentrifuge tubes were placed on ice immediately after collection. Blood samples were centrifuged at 3000 rpm for 15 min at 4°C. Extracted plasma was aliquoted into Eppendorf tubes and stored at -80°C until further analysis. Leptin ELISA Analysis Blood leptin concentrations were analyzed via PeproTech Murine Leptin ABTS ELISA Development Kit (Thermofisher #900-K76) at room temperature, following manufacturer’s instructions. Buffers and solutions were provided via PeproTech ABTS ELISA Buffer Kit (Thermofisher #900-K00). ELISA plates were pre-coated with leptin capture antibody overnight, prior to assay initiation. Pre-coated plates were washed and blocked with kit provided reagents for 1 h. Mouse serum was diluted in 1:40 ratio with kit provided diluent, and 100 µL were added to wells in triplicate and incubated for 2 h in the dark. Plates were then washed, and detection antibody was added and incubated for 2 h. After incubation, plates were again washed, and Avidin-HRP Conjugate was added and incubated for 3 min. Plates for washed for a final time and ABTS liquid substrate was added to all wells. After 25 min incubation, color development was monitored and measured the spectrophotometer at 405 nm with 650 nm correction.

### Tissue Harvest and RNA and Protein Sample Processing

All isolated tissue samples were freshly harvested and quickly snap-frozen in liquid nitrogen and stored in -80°C until they were processed for downstream analyses.

To isolate total RNA, individual tissues were bead homogenized in TRIzol using the Qiagen TissueLyser II and total RNA was extracted using the TRIzol reagent method according to the manufacturer’s instruction. Total RNA (2ug) was reverse transcribed to cDNA using the ThermoFisher High-Capacity cDNA Reverse Transcription Kit. RT-qPCR was performed using SYBR green and the Bio-Rad CFX384 Real-Time PCR System.

To isolate total protein, individual tissues were bead homogenized in 2% SDS lysis buffer supplemented with protease and phosphatase inhibitors using the Qiagen TissueLyser II at 4°C and centrifuged at max speed (∼21,000g) for 15 min at 4°C to remove debris. All fat tissue samples underwent at least 3 additional centrifuge spins to remove remaining fat layer. Brown adipocyte samples were rotated in either 2% SDS or RIPA buffer for 1 h at 4°C, sonicated on high for 10 cycles (30 sec on, 30 sec off; 10 min total) at 4°C, and then centrifuged at max speed (∼21,000g) for 15 min at 4°C. Protein concentrations for both types of samples were determined by Pierce bicinchoninic assay (BCA) and the Gen 5 v2.01.14 software. Protein samples were resolved on 12% SDS-PAGE gels and transferred to polyvinylidene fluoride (PVDF) membranes. Blots were probed with target antibodies and visualized using the FluorChem imaging system. Images were quantified with densitometry using Fuji (ImageJ2 v2.9.0).

### Cellular Respiration Assay

Oxygen consumption rate (OCR) was measured by Seahorse XFe24 analyzer (Agilent). For mitochondrial stress test cellular respirometry analyses, 2.0x10^4^ primary brown adipocytes were seeded and differentiated for 5 days (as previously described) in 2% gelatin coated XFe24 cell cultures plates. On the day of the assay (days 5-7 of differentiation), cells were washed and switched to unbuffered DMEM supplemented with 4.5g/L glucose, 4mM glutamine, 100mM sodium pyruvate and 2% of fatty acid-free BSA, pH 7.4. Drug concentrations in well (∼500µL starting volume): CL 316,243 (5µM), Oligomycin (4.5µM), DNP (1mM), Rotenone/Antimycin A (3.75µM each). For CL treatment measurements: 1 min mix, 0 min wait, 2 mins measure per cycle for 10 cycles. For all other measurements (basal, oligo, DNP, Rot/ AA): 1 min mix, 2 mins wait, 3 mins measure per cycle for 3 cycles. To measure extracellular acidification rates (ECAR) cell culture and differentiation protocols were followed as previously described. On the day of the assay (days 5-7 of differentiation), cells were washed and switched to unbuffered DMEM supplemented with 2mM glutamine, pH 7.4. Drug concentrations in well (∼500µL starting volume): Glucose (11mM), Oligomycin (4.5µM), 2-D-Glucose (50mM). All measurements: 30 sec mix, 1:30 mins wait, 2:30 mins measure per cycle for 3 cycles. Respirometry data was collected using the Agilent Wave software v2.6.1 and exported to GraphPad Prism v10.

### Body Composition

Lean, fat, and free fluid masses were measured by nuclear magnetic resonance (NMR) via the Minispec LF65 Body Composition Mice Analyzer (Bruker, Karlsruhe, Germany) as described previously (57).

### Whole Brown Adipose Tissue Proteomics

To analyze the whole cell proteome, brown adipose fat pads from male wild-type mice were isolated and lysed in 2% SDS solution supplemented with protease and phosphatase inhibitors. Sample lysates were quantified by bicinchoninic protein assay and delivered to the Biotechnology Resource Center (BRC) at Cornell University for Tandem Mass-Tagged (TMT) shotgun-based quantitative proteomics. Briefly, proteins were denatured, reduced, cysteine blocked, and digested using the S-trap approach. The resulting tryptic peptides were TMT-labeled and pooled. The labeled peptides were then fractionated by high pH reverse phase chromatography by the Ultimate 3000 MDLC platform into 10 fractions. Then samples were subjected to nanoLC-MS/MS analysis using a reverse phase HPLC separation and NanoLC RP coupled with an Orbitrap Eclipse mass spectrometer (Thermo Scientific) equipped with a nano ion source. Ion quantification and proteomic database searches were conducted using the Proteome Discoverer 2.4 software against the mouse database. All MS and MS/MS raw spectra were processed using Proteome Discoverer 2.4 (PD 2.4, Thermo) for reporter ion quantitation analysis.

BAT protein samples from AQP1-Flox and KO mice on 60% HFD were processed in 2% SDS as previously described and sent to Weill Cornell Medicine Proteomics and Metabolomics Core Facility 16plex TMT for labeled proteomic analysis. Briefly, samples were precipitated by acetone and digested by trypsin. Desalted peptides were labeled and fractionated by RPLC into 12 fractions. The peptides of each fraction were desalted by C18 tips before they were analyzed by LC-MS. Each sample was analyzed using data dependent acquisition (DDA) method. Data was then searched against a Uniprot mouse protein database using the MaxQuant software.

#### Enzymatic digestion

Proteins were precipitated from lysate using 80% acetone, reduced with 5mM dithiothreitol (DTT) and then alkylated with 15mM iodoacetamide in the dark. The samples were digested with sequencing grade trypsin (Promega) overnight at 37°C. Digests were acidified by the addition of 10% trifluoroacetic acid (TFA) to 0.5% final concentration and the peptides were desalted on tC18 Sep-Pak cartridges (Waters) and dried in a centrifugal evaporator.

#### TMT-labeling

Peptides were resuspended in 0.2M HEPES buffer, pH 8.5. TMT amino reactive reagents (Thermo Fisher Scientific) were resuspended in 100 µl of anhydrous acetonitrile (ACN) and all 100 µl of each reagent was added to each sample and mixed briefly on a vortexer. Reactions were allowed to proceed at room temperature for 1 h, and then quenched by the addition of 200 µl of 5% hydroxylamine for 15 min and then acidified by the addition of 400 µl 100% FA. All TMT channels were combined in equal ratios and desalted.

#### Peptide pre-fractionation by high pH reverse phase chromatography

The TMT-labeled peptides were fractionated using high-pH reverse-phase HPLC. Separations were performed using an Agilent 1260 pump and a 4.6 mm × 250 mm XBridge C18, 5µm column (Waters) with a 50 min gradient from 18% to 38% buffer B (90% ACN, 10mM NH4HCO3, pH 8) at a flow rate of 0.8 mL/min. In total, 12 fractions were obtained.

#### LC-MS/MS Analysis of TMT samples

A Thermo Fisher Scientific EASY-nLC 1200 coupled on-line to a Fusion Lumos mass spectrometer (Thermo Fisher Scientific) was used. Buffer A (0.1% FA in water) and buffer B (0.1% FA in 80% ACN) were used as mobile phases for gradient separation. A 75 µm x 15 cm chromatography column (ReproSil-Pur C18-AQ, 3 µm, Dr. Maisch GmbH, German) was packed in-house for peptide separation. Peptides were separated with a gradient of 10–40% buffer B over 110 min, 40%-80% B over 10 min at a flow rate of 300 nL/min. The Fusion Lumos mass spectrometer was operated in data dependent mode. Full MS scans were acquired in the Orbitrap mass analyzer over a range of 400-1500 m/z with resolution 60,000 at m/z 200. The top 15 most abundant precursors with charge states between 2 and 6 were selected with an isolation window of 0.7 Thomson by the quadrupole and fragmented by higher-energy collisional dissociation with normalized collision energy of 40. MS/MS scans were acquired in the Orbitrap mass analyzer with resolution 30,000 at m/z 200. The automatic gain control target value was 1e6 for full scans and 5e4 for MS/MS scans respectively, and the maximum ion injection time was 100 ms for MS scans and 54 ms for MS/MS scans.

The raw files were processed using the MaxQuant computational proteomics platform (version 2.4.2.0) for protein identification. The fragmentation spectra were used to search the UniProt mouse protein database (downloaded on 09/17/2021). TMT on peptide N-term / lysine, oxidation of methionine and protein N-terminal acetylation were used as variable modifications for database searching. Cysteine carbamidomethylation was used as a fixed modification. The precursor and fragment mass tolerances were set to 7 and 20 ppm, respectively. Both peptide and protein identifications were filtered at 1% false discovery rate based on decoy search using a database with the protein sequences reversed. Gene ontology analysis was performed using the Gene Ontology Consortium knowledgebase (geneontology.org) (74, 75).

### Glucose Uptake Rate

AQP1-Flox and KO primary murine brown adipocytes (2.0x10^4^cells/well) were seeded and differentiated for 5 days (as previously described above) in 2% gelatin coated white, opaque, flat-bottom 96 well plates. On the day before the assay, cells were treated with maintenance media without serum and insulin for 24 h. The day of the assay, media was carefully replaced with buffered DMEM (2.438g/L sodium bicarbonate) without serum or glucose and incubated in the cell culture incubator (5% CO_2_, 37°C) for 30 min. Cell culture media was then replaced with 1µM insulin in DMEM without serum or glucose and incubated in the cell culture incubator for 1 h. The glucose uptake rate was then determined using the Glucose Uptake-Glo Assay kit (Promega) according to manufacturer’s instructions. All controls and groups were plated in duplicate. Briefly, a 2DG6P standard curve from 0-50µM was diluted in PBS and the 2DG6P Detection was incubated at 25°C in the dark during the 1 h insulin treatment. After the 1 h insulin treatment, 1mM 2DG was added to all wells (excluding basal control duplicates) and incubated at 25°C for 5 min. 25uL of Stop buffer was added to each well followed by 25µL of Neutralization buffer with brief plate shaking to mix after each step. 100µL of 2DG6P Detection Reagent was added to each well and after briefly shaking the plate to mix, the plate was incubated in the dark at 25°C for 1 h. Luminescence signal was recorded using a 0.3-1 second integration on the EnVision Nexus Multimode Microplate Reader (Revvity).

### Quantification and Statistical Analysis

Statistical analyses were performed using GraphPad Prism 10 software in consultation with the Cornell Statistical Consulting Unit (CSCU). Specific statistical tests are indicated in each Fig.. Most data are represented as the mean ± S.E.M. unless otherwise indicated. Both unpaired two-tailed Student’s T-test and two-way ANOVA were used with Bonferroni post hoc analysis.

All *ex vivo* studies used independently derived primary brown adipocytes and had at least two biological replicates conducted. For animal experiments, an n=6-12 per group for an alpha value of 0.05 and an effect size of >10-20% respectively was determined to yield significance based on previous studies.

## Author Contributions

Conceptualization-J.J.B. and C.M.C.

Methodology-J.J.B., C.M.C., C.J.B., P.T.

Investigation-C.M.C., C.J.B., P.T., K.E., M.L., Y.Q., Y.L., N.B., M.S.

Validation-C.M.C., C.J.B., M.L., N.B., M.S.,

Formal Analysis-J.J.B., C.M.C. Writing-J.J.B., C.M.C.

Review and Editing-J.J.B., C.M.C., C.J.B., P.T., K.E.

Visualization-J.J.B., C.M.C. Supervision-J.J.B.

## Acknowledgements

Dr. Olivier Devuyst from the Institute of Physiology at the University of Zurich for sharing the AQP1-Flox mouse and Sebastien Druart and Delphine De Mulder from UCLouvain for genotyping information and shipment of AQP1-Flox mice.

Aric Kraft and Mike Kacergis from Sable Systems for their expertise regarding Promethion Metabolic Cage analysis and management.

Dr. Guoan Zhang from Weill Cornell Medicine Proteomics and Metabolomics Core Facility for conducting proteomics experiment for HFD AQP1-Flox and KO mice.

Dr. Shannon Reilly from Weill Cornell Medicine for technical advice.

Cornell CARE staff for the high quality and careful management of animals used in the study. Cornell BRC for conducting the proteomics from WT mice.

## Supporting Information

**Fig. S1.**
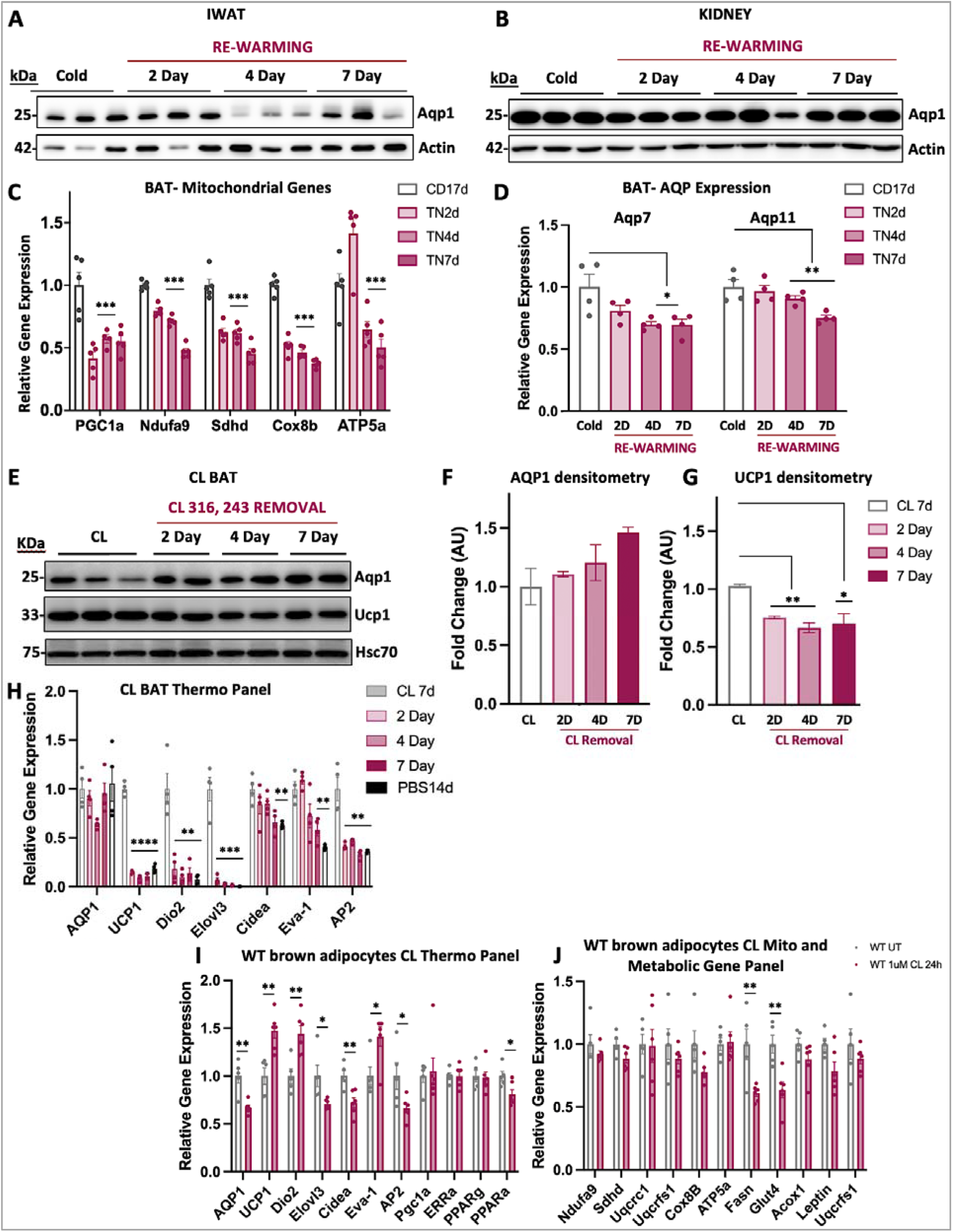
Characterization of thermogenic biomarker gene expression in BAT, iWAT, and kidney harvested from WT male mice under NST active and silencing conditions. (A) Representative immunoblot of iWAT isolated from WT male mice exposed to CD 7d, compared to mice exposed to CD for at least 7 days followed by a re-warming period of 2, 4, or 7 days in TN (n=5). (B) Representative immunoblot of kidney isolated from WT male mice exposed to CD 7d, compared to mice exposed to CD for at least 7 days followed by a re-warming period of 2, 4, or 7 days in TN (n=5). (C) Relative mRNA expression of mitochondrial biomarkers from BAT isolated from mice exposed to CD 7d or CD 7d followed by re-warming period (n=5). (D) Relative mRNA expression of other relevant aquaporins from BAT isolated from mice exposed to CD 7d or CD 7d followed by re-warming period (n=5). (E) Representative immunoblot of BAT isolated from WT male mice exposed to injected with 1mg/kg CL 316,243 (CL) daily for 14 days compared to mice injected with CL for at least 7 days followed by a CL removal (i.e. “re-warming” period) of PBS injections for 2, 4, or 7 days in TN. (n=5). (F) Densitometry quantification of AQP1 protein expression normalized to loading control HSC70 from S1E immunoblot under CL or CL followed by PBS injection conditions at TN. (G) Densitometry quantification of UCP1 protein expression normalized to loading control HSC70 from S1E immunoblot under CL or CL followed by PBS injection conditions at TN. (H) Relative mRNA expression of thermogenic biomarkers from BAT isolated from mice under CL or CL followed by PBS injection conditions at TN (n=5). (I) Relative mRNA expression of thermogenic biomarkers from WT primary brown adipocytes treated with PBS vehicle or 1uM CL for 24h. Groups are WT UT (n=5) or WT 1uM CL (n=6). (J) Relative mRNA expression of mitochondrial or metabolism biomarkers from WT primary brown adipocytes treated with PBS vehicle or 1uM CL for 24h. Groups are WT UT (n=5) or WT 1uM CL (n=6). Densitometry and qPCR data represented as mean + SEM. Significance is denoted as *p<0.05, **p<0.01, ***p<0.001 by Student’s t test.

**Fig. S2.**
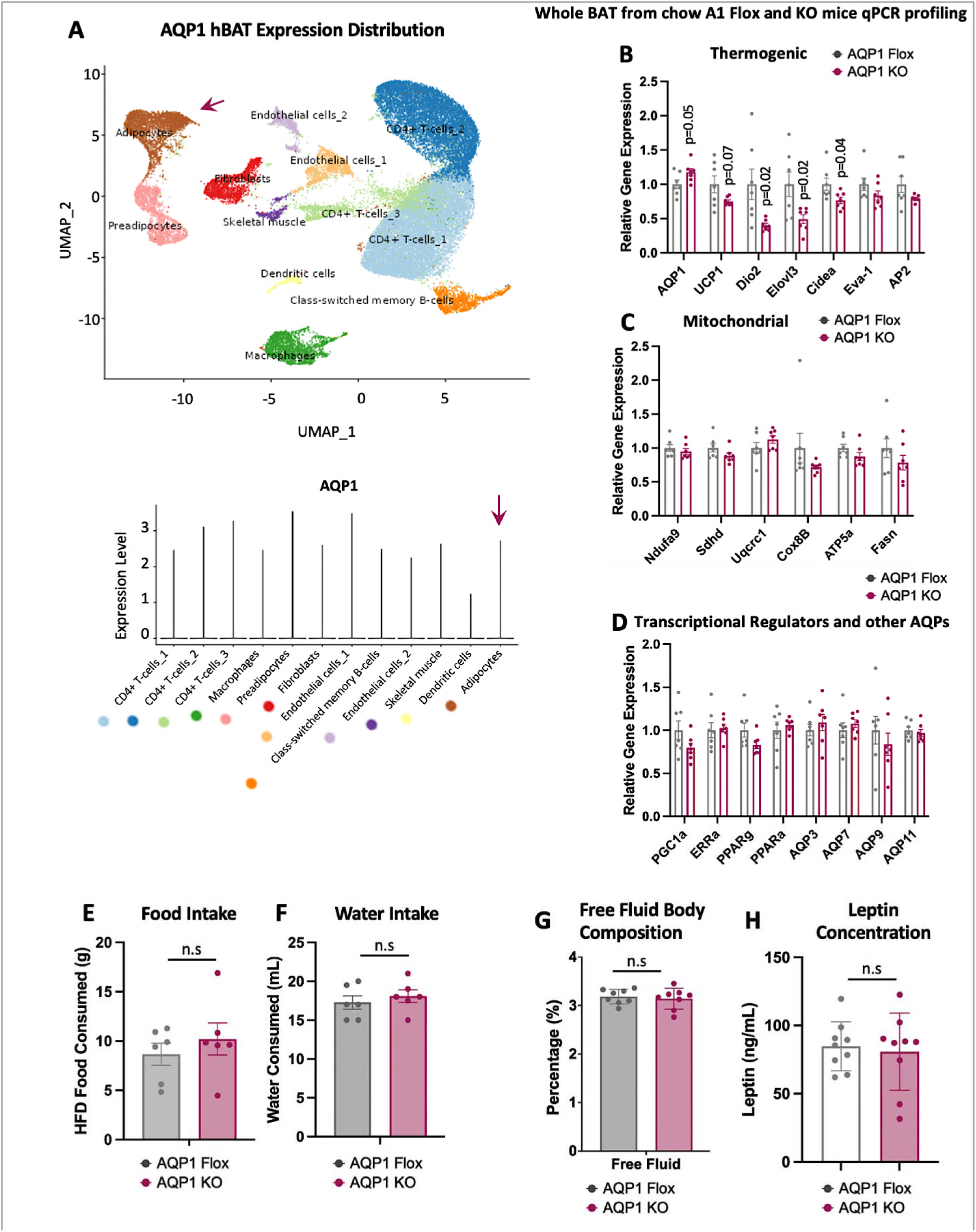
Characterization of murine AQP1-KO mice in chow and HFD dietary conditions. (A) AQP1 gene expression from previously published snRNA-seq database (43) shown as uniform manifold approximation and projection (UMAP) plot for different cell populations in human BAT coupled with violin plot for AQP1 expression in different cell types. (B-D) Relative mRNA expression in whole BAT from chow-fed AQP1-Flox and AQP1-KO male mice (n=7/group) analyzing (B) thermogenic markers, (C) mitochondrial markers, and (D) transcriptional regulators and other aquaporins (AQPs). (E) Food intake measured after one week of single housing AQP1-Flox or KO mice in TN conditions after 24 weeks of HFD (n=6). (F) Water intake measured after one week of single housing AQP1-Flox or KO mice in TN conditions after 24 weeks of HFD (n=6). (G) Body composition measured by nuclear magnetic resonance (NMR) to measure free fluid percentage (n=8). (H) Plasma leptin concentration from AQP1-Flox and AQP1-KO mice after 25 weeks of HFD (n=9). All bar graphs are represented as mean + SEM. Significance is denoted as *p<0.05, **p<0.01, ***p<0.001 by Student’s t test.

**Fig. S3.**
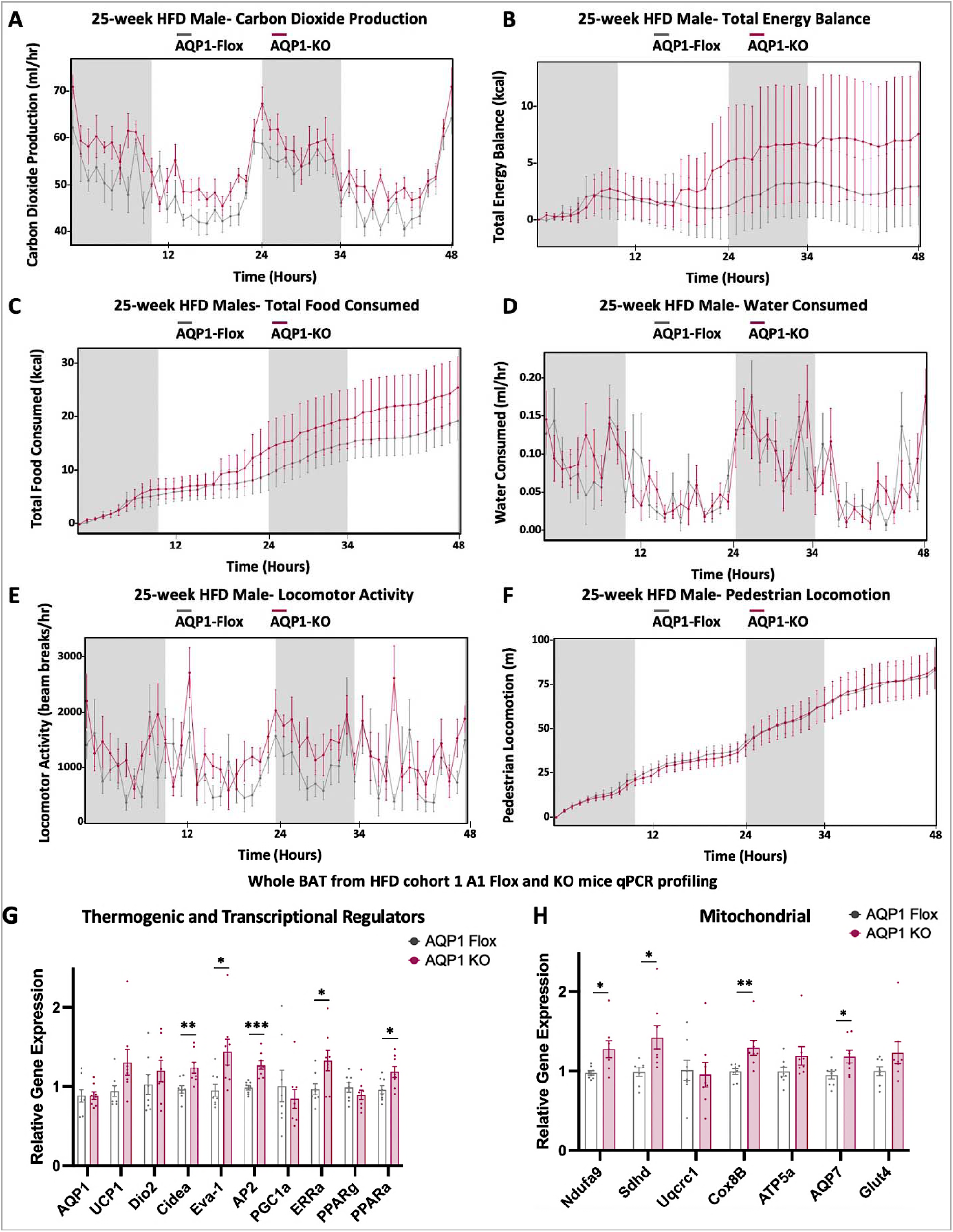
*In vivo* characterization of AQP1-Flox and KO mice metabolic parameters after 24 weeks of HFD in TN conditions. (A-F) Metabolic data obtained from Promethion metabolic cages of AQP1-Flox and AQP1-KO male mice single-housed at 30°C for a 48h period after 24 weeks of HFD (n=6). (A) Carbon dioxide production. (B) Total energy balance. (C) Total food consumption. (D) Water consumption. (E) Locomotion Activity. (F) Pedestrian Locomotion. (G-H) Relative mRNA expression of biomarkers in BAT from AQP1-Flox and AQP1-KO male mice after 25 weeks of HFD at 30°C. (G) Thermogenic and transcriptional regulators. (H) Mitochondrial markers. Relative mRNA expression data represented as mean + SEM. Significance is denoted as *p<0.05, **p<0.01, ***p<0.001 by Student’s t test.

**Table S1.**
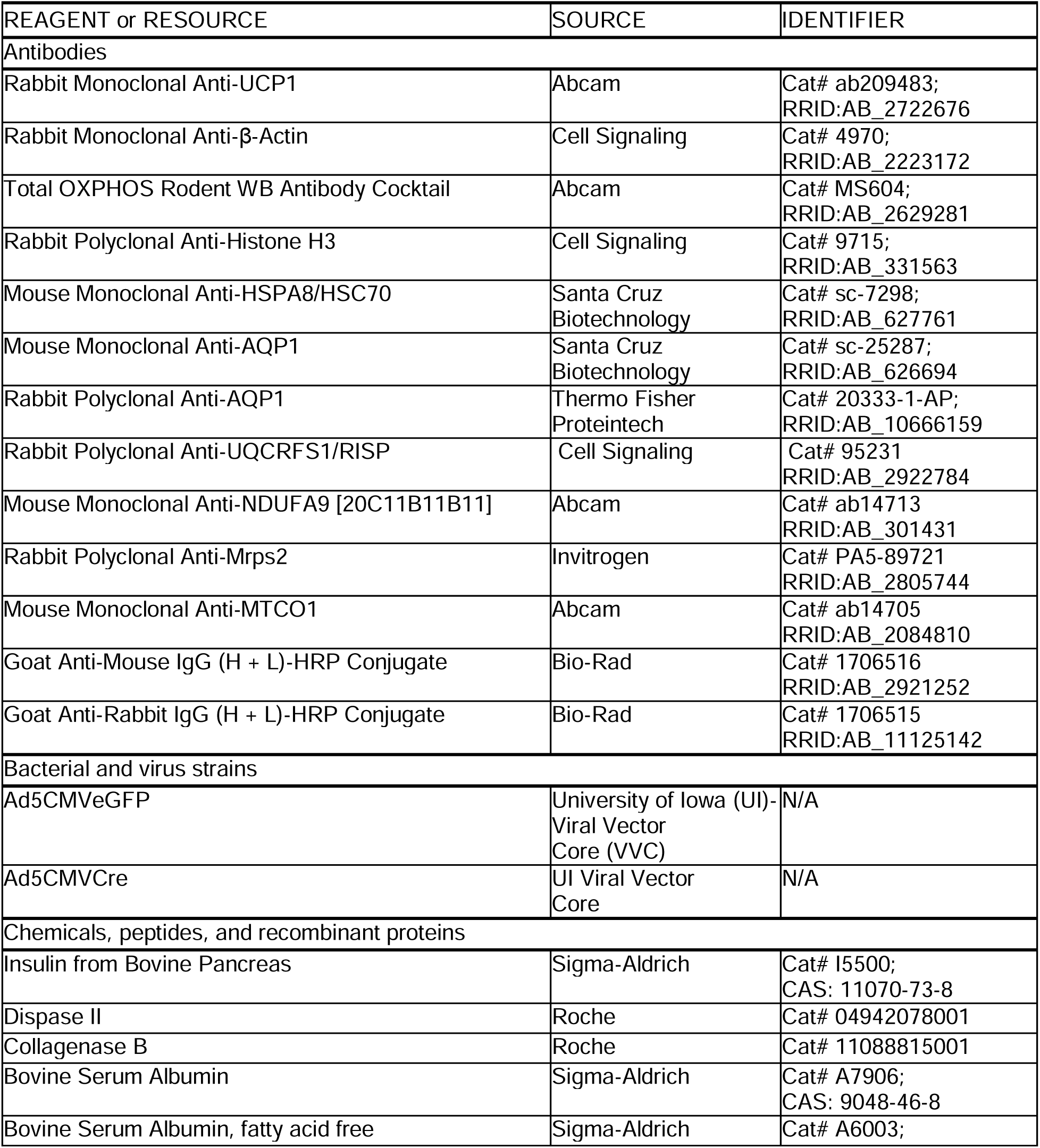
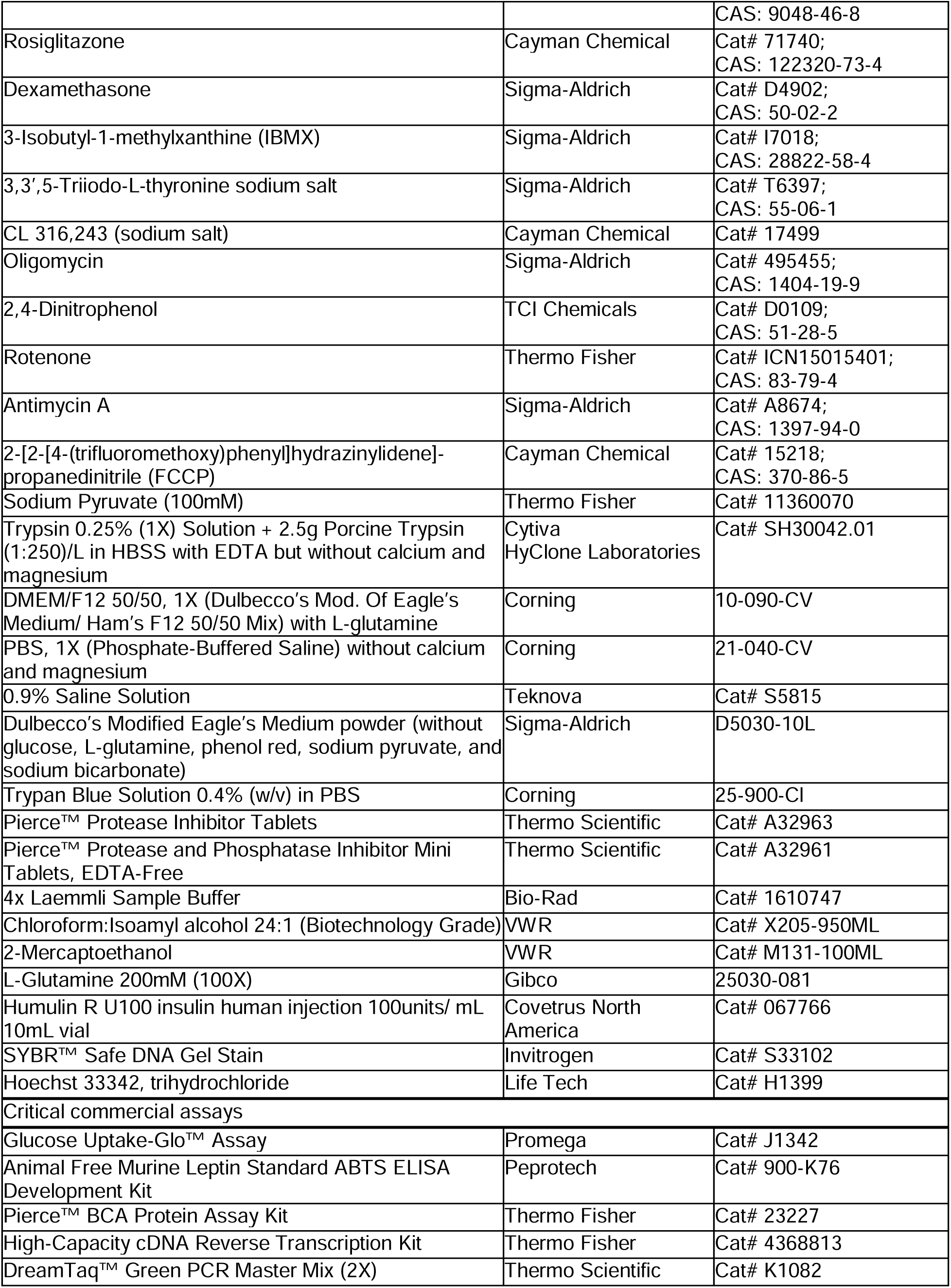
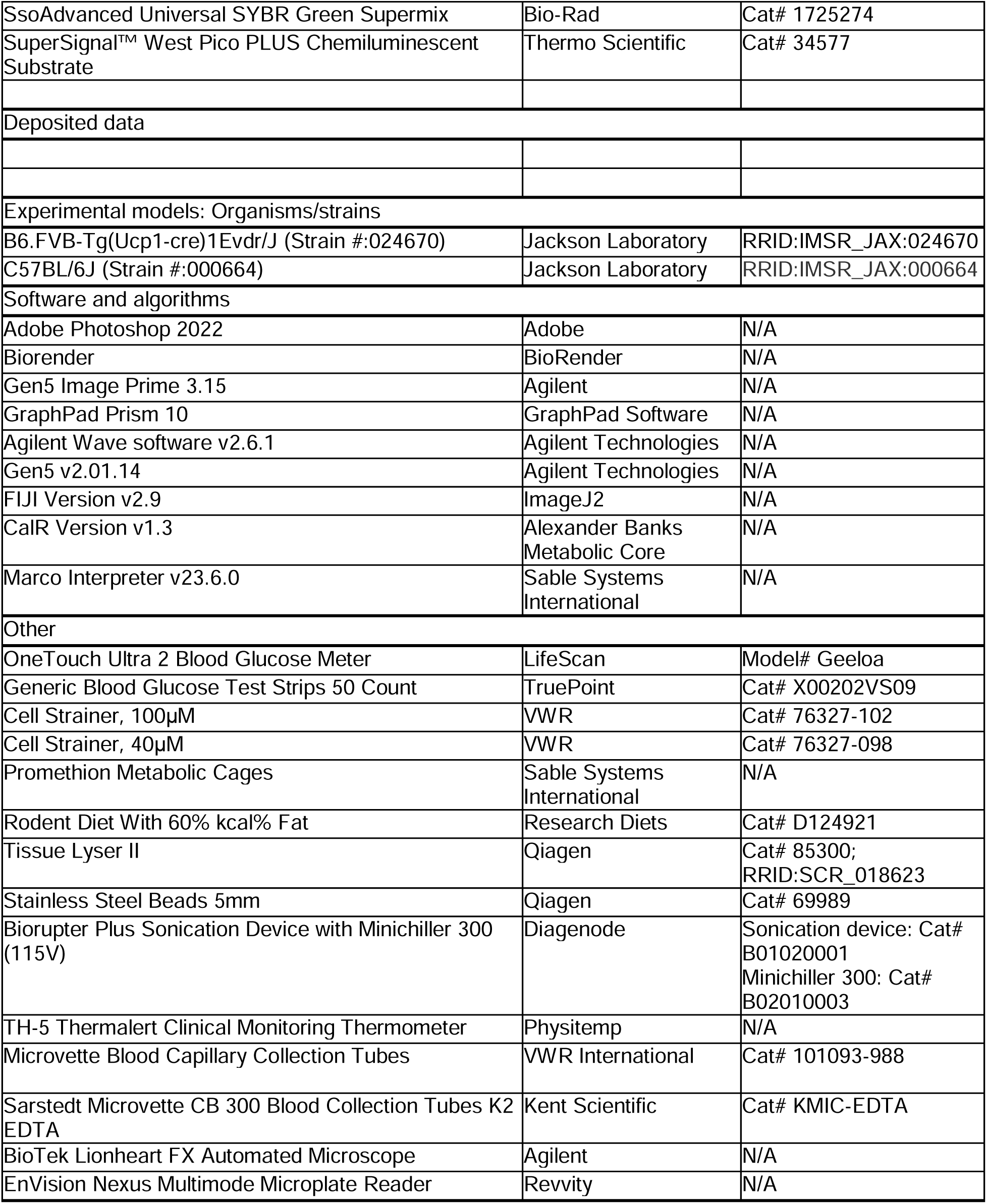
Key resources table.

**Table S2.**
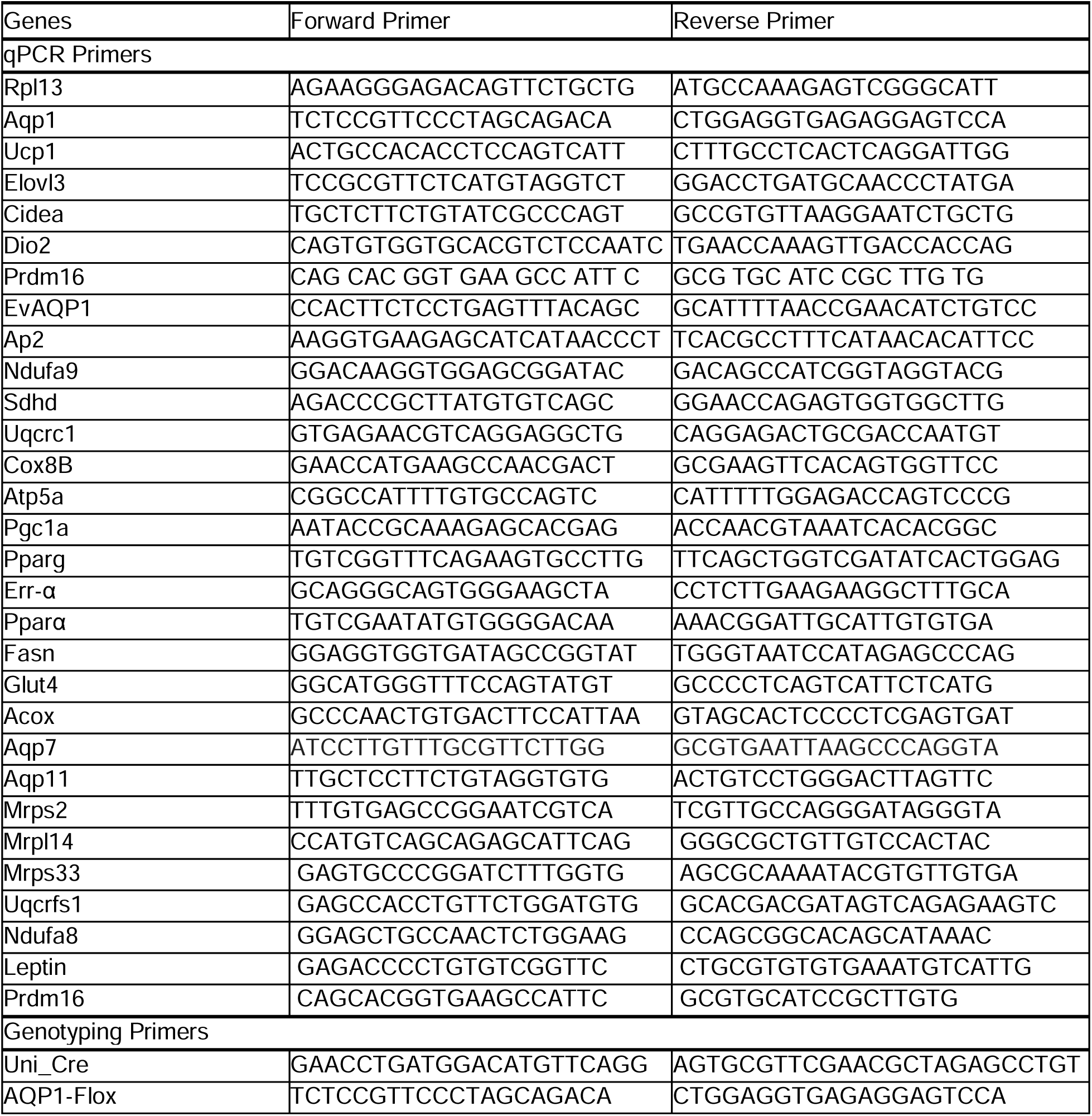
Primer List.

